# Phylogenetic scale in ecology and evolution

**DOI:** 10.1101/063560

**Authors:** Catherine H. Graham, David Storch, Antonin Machac

**Affiliations:** Department of Ecology and Evolution, 650 Life Sciences Bldg, Stony Brook University, Stony Brook, NY 11794, USA; Swiss Federal Research Center (WSL) Birmensdorf, Switzerland; Department of Ecology, Vinicna 7, 12844 Prague 2, Czech Republic; Center for Theoretical Study, Jilska 1, 11000 Prague 1, Czech Republic; Center for Macroecology, Evolution, and Climate, Natural History Museum of Denmark, Universitetsparken 15, DK 2100 Copenhagen, Denmark

**Keywords:** biogeography, community structure, diversification, domains of scale, extent, grain, macroecology, macroevolution, spatial scale

## Abstract

**Aim:** Many important patterns and processes vary across the phylogeny and depend on phylogenetic scale. Yet, phylogenetic scale has never been formally conceptualized and its potential remains largely unexplored. Here, we formalize the concept of phylogenetic scale, review how phylogenetic scale has been considered across multiple fields, and provide practical guidelines for the use of phylogenetic scale to address a range of biological questions.

**Methods:** We summarize how phylogenetic scale has been treated in macroevolution, community ecology, biogeography, and macroecology, illustrating how it can inform, and possibly resolve, some of the longstanding controversies in these fields. To promote the concept empirically, we define phylogenetic grain and extent, scale-dependence, scaling, and the domains of phylogenetic scale. We illustrate how existing phylogenetic data and statistical tools can be employed to investigate the effects of scale on a variety of well-known patterns and processes, including diversification rates, community structure, niche conservatism, or species-abundance distributions.

**Main conclusions:** Explicit consideration of phylogenetic scale can provide new and more complete insight into many longstanding questions across multiple fields (macroevolution, community ecology, biogeography, macroevolution). Building on the existing resources and isolated efforts across fields, future research centered on phylogenetic scale might enrich our understanding of the processes that together, but over different scales, shape the diversity of life.

## Introduction

Numerous patterns in ecology and evolution vary across the phylogenetic hierarchy (Fig. 1). Species diversity declines with latitude across orders and classes, but not necessarily across their constituent families and genera (Marquet *et al*., 2004; Buckley *et al*., 2010). Phylogenetic delimitation of species pools influences our inferences about the processes that form local communities (Cavender-Bares *et al*., 2009; Chalmandrier *et al*., 2013). Many other, similar examples further illustrate that patterns in ecology and evolution often depend on phylogenetic scale (Fig. 1). Yet, unlike the extensively developed concepts of spatial and temporal scale, where scale-dependence in the patterns and processes has long been acknowledged (Wiens, 1989), the importance of phylogenetic scale has only recently begun to be recognized. Here, we formalize and develop the concept of phylogenetic scale, summarize how it has been considered across fields, provide empirical guidelines for the treatment of phylogenetic scale, and suggest further research directions.

**Figure 1.**
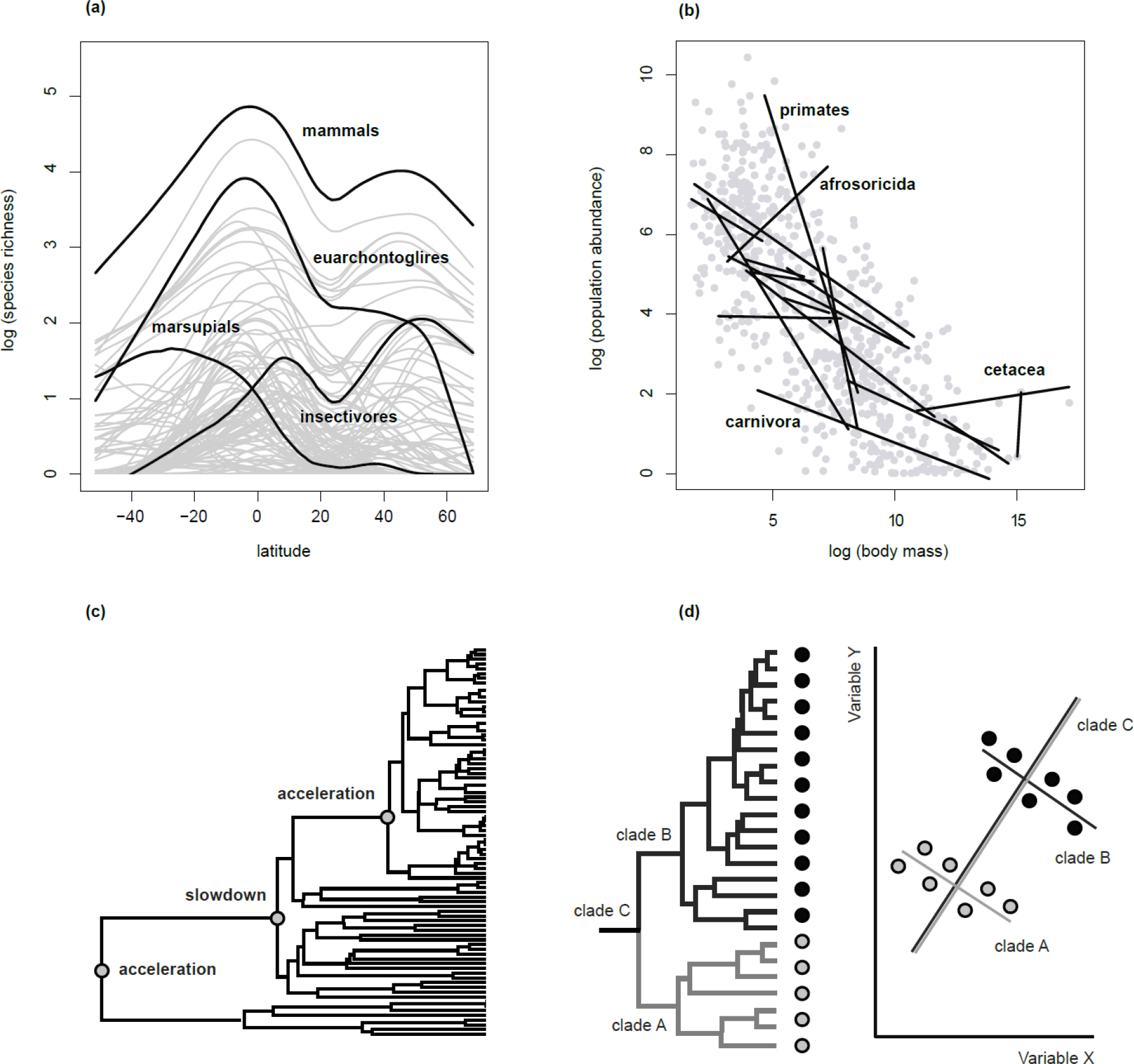
Examples of patterns that vary across phylogenetic scales. (a) The latitudinal diversity gradient. Mammal diversity decreases with latitude across large clades but many other patterns emerge across small clades, including inverse ones (selected clades depicted in black). (b) The dependence of population abundance on body mass. The dependence is negative across large phylogenetic scales (mammals depicted in grey) but varies substantially across small scales (selected orders depicted in black). (c) Diversification dynamics. Slowdowns detected over some phylogenetic scales might be accompanied by accelerations over both larger and smaller scales. (d) Statistical correlations. Even though the depicted variables are positively correlated within each of the two subclades, the correlation becomes negative when the subclades are studied together. The data were taken from the IUCN Database (IUCN 2017) and PanTHERIA (Jones *et al*., 2009).

Scale is a concept derived from the fact that entities, such as clades, can be ordered within a hierarchy, which in this essay is represented by the phylogenetic hierarchy. Phylogenetic scale describes the position of a clade within this hierarchy, relative to other clades, and can be measured in terms of tree depth, taxonomic rank, or clade age. In some cases, by analogy with spatial scale (Wiens, 1989), we can distinguish between phylogenetic extent and phylogenetic grain (Box 1). Phylogenetic extent and grain are defined in relation to each other, where extent refers to the entire phylogeny under study, while grain refers to those parts of the phylogeny (e.g. nodes, subclades) that represent the elementary unit of analysis. In community ecology, for example, analyses based on large phylogenetic extents, such as angiosperms, typically suggest that community composition has been shaped by environmental filters while analyses of narrowly defined extents (e.g. white oaks), often implicate a suite of additional mechanisms (such as competition, mutualisms, dispersal limitation) (Cavender-Bares *et al*., 2009). Researchers working within the same phylogenetic extent (e.g. families of birds), but across multiple grains (clades within different depths of the family phylogeny), found that clades with deep divergence times show higher sympatry than younger clades, as expected under the model of allopatric speciation followed by secondary sympatry (Barraclough & Vogler, 2000). These examples illustrate that phylogenetic extent and grain might be useful conceptual tools facilitating a thorough examination of various patterns across the phylogeny. However, there are instances when the distinction between extent and grain might not be necessary or meaningful (Box 1), in which case we use the general term phylogenetic scale.

The concept of phylogenetic scale seems particularly pertinent, given the growing body of research to explore the increasingly accurate and ever more complete phylogenetic data (Table 1, Table 2). Yet, few studies have systematically investigated how different patterns and processes change with phylogenetic scale (e.g. niche conservatism, community structure, diversification rate, extinction risk), or used phylogenetic scale to identify the “laws of ecology” (i.e. patterns that hold across scales, such as species-abundance distributions, or latitudinal gradients).We contend that the full potential of the phylogenetic data, together with that of the methods currently available to analyze these data (Table 1, Table 2), have not yet been fully realized. More rigorous treatment of phylogenetic scale may consequently generate a more in-depth understanding of biological patterns and processes, similar to that previously produced by temporal and spatial scale (Wiens, 1989).

Here, we overview the variety of ways in which different fields of ecology and evolution have implicitly or explicitly considered phylogenetic scale. Then, we propose a conceptual framework that might provide the common ground for cross-field discussion. In particular, we define and formalize the concept of phylogenetic scale, introduce phylogenetic grain and extent, and define further potentially useful concepts, such as scale-dependence, phylogenetic scaling and domains of scale (Box 2). We also provide practical guidelines for the treatment of phylogenetic scale across empirical studies, using phylogenetic data and statistical methods currently available. We hope our effort will inspire further debate, draw more focused attention to the subject, and advance the notion of phylogenetic scale in ecology and evolution.

### BOX 1: The concept of phylogenetic scale

The concept of scale is based on the fact that some entities can be ordered, or placed on a scale (*scala* means *ladder* in Latin). For example, continents contain biomes, ecoregions, and localities, giving rise to a nested spatial hierarchy. Similarly, large clades contain small clades, creating phylogenetic hierarchy which defines phylogenetic scale. Phylogenetic scale refers to the position within this hierarchy.

Different measures might be used to represent phylogenetic scale, depending on the biological rationale of the study and the question addressed. Standardized measures are particularly pertinent when the studied clades are mutually non-nested, such that we cannot readily refer to the phylogenetic hierarchy. In these cases, taxonomy has been traditionally used to position nested as well non-nested clades along the scale continuum. Yet, taxonomy is rarely the best measure of phylogenetic scale because its ranks, especially for unrelated taxa, might not be fully comparable (e.g. genera in mammals are not comparable to genera in insects). For this reason, other standardized measures, such as tree depth, clade age, clade size, or the degree of molecular and phenotypic divergence, might be more appropriate for most questions. Even these measures, however, have some limitations (e.g. clades of the same age might not be comparable in terms of their diversification rates, niche conservatism, etc.), but these limitations apply to the measures of spatial scale as well. Spatial grains of standardized sizes, for example, might not ensure comparability across species with dramatically different home range sizes (Wiens, 1989). Consequently, there is not a single all-purpose measure for phylogenetic scale. Instead, the choice of the most suitable measure will typically stem from the biological properties of the organismal system studied (e.g. body size, generation time, rates of morphological evolution) and the attribute we wish to evaluate (e.g. diversification, niche conservatism, phenotypic divergence) (Box 2).

The principal difference between phylogenetic and temporal scale is that the latter ignores phylogenetic hierarchy. Even though time might serve as a practical surrogate for phylogenetic scale in some cases (Jablonski, 2007), it typically will not fully capture the variation in the attribute of interest (e.g. diversification, niche conservatism, trait divergence) across the entire phylogeny. For example, the same trait might evolve at different rates even across closely related clades (e.g. due to clade-specific selection regimes), such that the same temporal scale becomes associated with very different rates of trait evolution, and by extension, with different degrees of niche conservatism. In this case, the degree of trait divergence might serve as a more suitable (time-independent) measure of phylogenetic scale, which allows delimitation of mutually comparable clades in terms of their niche conservatism. The concept of phylogenetic scale may therefore encourage a more accurate way of thinking about patterns and processes across the phylogenetic hierarchy.

Phylogenetic scale can be further defined in terms of phylogenetic grain and phylogenetic extent. In spatial scale, grain typically refers to the area of the elementary unit of analysis (e.g. ecoregions or grid cells within a continent) while extent refers to the total area analyzed (e.g. continent). Similarly, phylogenetic grain refers to the elementary unit of analysis, defined in terms of tree depth, taxonomic rank, clade age, or clade size, and phylogenetic extent refers to the total phylogeny that would encompass all the units analyzed (= all grains). Extent and grain are defined in relation to each other, such that the grain from one study can act as an extent in another study, and vice versa. Importantly, the distinction between extent and grain might be required for some questions, but might not be useful for others. For example, rates of trait evolution are associated exclusively with the clade under analysis, irrespective of other clades in the phylogeny analyzed. In this case, it is not meaningful to distinguish between the extent and grain of trait conservatism.

**Figure:** Comparison of spatial and phylogenetic scale. Both scales describe a hierarchy. Spatial scale is typically defined in terms of geographic area. Phylogenetic scale can be defined in terms of clade age, tree depth, or taxonomic ranks. In both of the depicted examples, grain B represents a larger unit of the analysis that contains multiple smaller units of grain A. Extent encompasses all the units analyzed.

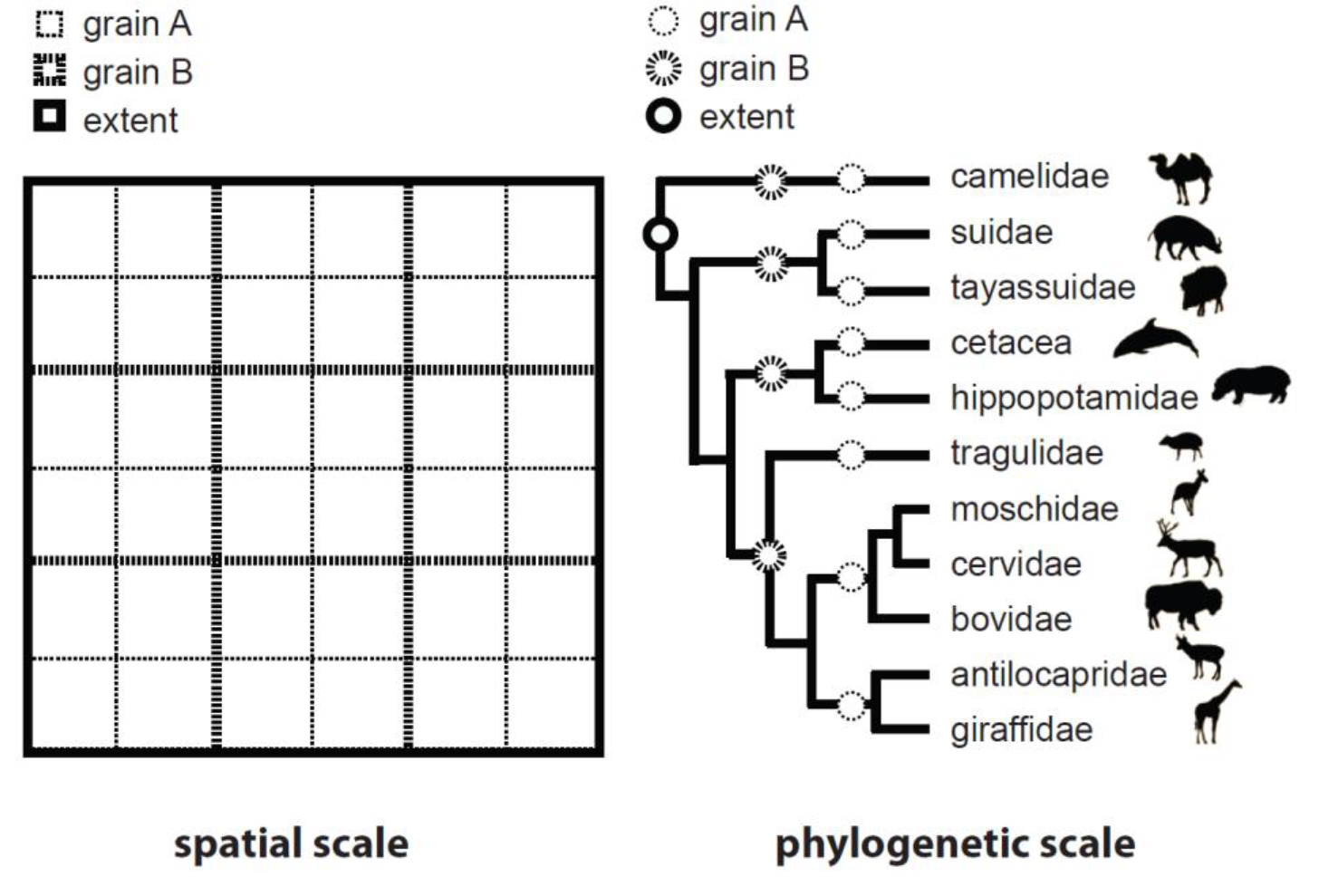

## Phylogenetic scale in ecology and evolution

Different fields in ecology and evolution have considered the concept of phylogenetic scale to varying degrees, from acknowledging that patterns change across scales to explicit scale-based analyses. In this section, we describe research that has treated, implicitly or explicitly, phylogenetic scale and suggest how different fields might further benefit from this concept.

### Evolution and diversification

Evolutionary diversification (net outcome of speciation and extinction) and disparification (divergence of trait values within a clade) are known to vary across phylogenetic scales, but rarely studied in this context. Even though there are a suite of methods to explore diversification and disparification across the phylogeny (Alfaro *et al*., 2009; Ingram & Mahler, 2013; Rabosky, 2014) (Table 2), most studies report macroevolutionary patterns without systematically investigating how the patterns change with respect to phylogenetic scale. Such an examination across phylogenetic grains (different clades within the same phylogenetic extent) and extents (clades within different extents, such as genus, family, class, etc.) may produce a more complete picture of the analyzed patterns and help resolve some of the outstanding controversies in the field.

One such controversy revolves around the dynamics of diversity and diversification. It has been debated whether the dynamics are expansionary, such that regional and clade diversity accumulate constantly over time (Benton & Emerson, 2007; Harmon & Harrison, 2015), or whether the dynamics are ecologically limited, such that diversity tends toward an equilibrium (Rabosky & Hurlbert, 2015). Empirical evidence suggests that genera with dozens of species often expand in terms of their diversity (Benton & Emerson, 2007; Wiens, 2011; Harmon & Harrison, 2015). In contrast, higher taxa with thousands of species seem to be mostly saturated at their equilibrium diversity (Rabosky & Hurlbert, 2015); though there is evidence that some of the very large clades (e.g. birds, tetrapods) might diversify continuously as their select subclades saturate (Jetz *et al*., 2012; Hedges *et al*., 2015). Island radiations and fossil evidence also indicate that clades often expand, seemingly without bounds, during the initial phases of their diversification but eventually reach an equilibrium and saturate (Benton & Emerson, 2007; Glor, 2010). Even though some of the observed slowdowns might be caused by statistical artifacts, cryptic speciation, and other effects (Alizon *et al*., 2008; Moen & Morlon, 2014), it is possible that diversification tends to vary systematically across phylogenetic scales such that seemingly contradictory dynamics (e.g. expansionary and equilibrial) might be detected within the same phylogenetic tree (Fig. 1c) (Benton & Emerson, 2007; Jablonski, 2007; Jetz *et al*., 2012; Hedges *et al*., 2015). If this is the case, the debate as to whether the dynamics are expansionary or equilibrial should perhaps be reframed in terms of phylogenetic scale. For example, we could investigate the scales over which the different dynamics prevail, identify the ecological factors that determine the shifts between the dynamics, or study how the dynamics combine across nested clades of different ages and sizes to produce the emergent diversification dynamics, observed across the entire phylogeny (see Phylogenetic scale in practice) (Benton & Emerson, 2007; Jablonski, 2007; Machac *et al*., 2013).

Evolutionary disparification varies across the phylogeny, as well, because traits (phenotypic, physiological, behavioral, but also molecular) diverge at different rates and, therefore, are conserved over different phylogenetic scales (Blomberg *et al*., 2003; Harmon *et al*., 2010). Even though the rates of trait (or niche) evolution have been the subject of much research, clear generalizations about how they vary across phylogenetic scales have not yet emerged. In some cases, physiological traits that largely determine species distributions (e.g. frost tolerance) (Donoghue, 2008), might be conserved over extensive phylogenetic scales (e.g. at the family level) while habitat- and diet-related traits that mediate species coexistence locally are often labile and thus conserved over small scales (Blomberg *et al*., 2003). However, the opposite pattern has also been observed where physiological tolerances were conserved over small scales while habitat, diet, body size, and feeding method remained unchanged for most of a clade’s history (Price *et al*., 2014).

### Community ecology

Community ecology stands out as a field where the effects of phylogenetic scale have not only been recognized, but also extensively studied, thus illustrating the theoretical and empirical potential of the concept (Cavender-Bares *et al*., 2009; Munkemuller *et al*., 2014). Specifically, research across phylogenetic grains and extents has been used to disentangle the suite of processes that together shape community structure.

To study the phylogenetic structure of a community, researchers often calculate standardized metrics which can be classified with respect to the phylogenetic grain that they capture (Webb *et al*., 2002; Swenson, 2009, 2011; Mazel *et al*., 2016). The nearest taxon index (NTI), for example, targets the shallow parts of the phylogeny, or small phylogenetic grains, as it measures distances between closely related species within a community. The net relatedness index (NRI), in contrast, measures the distances between all species within a community, thus covering an inclusive range of grains, both small and large (Webb *et al*., 2002; Swenson, 2009). The same sensitivity to community structure at different phylogenetic grains holds for beta-diversity metrics (e.g. PhyloSor, UniFrac, and D_nn_ capture the shallow parts of the phylogeny) (Swenson, 2011). Combining metrics capturing different grains, Mazel *et al*. (2016) found evidence for recent diversification events in South America (phylogenetic clustering near the tips) but not in Africa (clustering near the root), suggesting that the faunas were assembled differently across the two continents (Mazel *et al*., 2016). Parmentier *et al*. (2014) investigated the structure of tree communities across a range of phylogenetic and spatial grains and concluded that environmental filtering shaped the communities at all but the smallest scales, where competition appeared to predominate (Parmentier *et al*., 2014).

Phylogenetic extent, too, can have significant effects on phylogenetic metrics of community structure. These metrics are often standardized with respect to null expectations, typically based on a species pool defined by the phylogenetic extent of the group under investigation (Cavender-Bares *et al*., 2009; Parra *et al*., 2010; Chalmandrier *et al*., 2013). Changing the phylogenetic extent of their analysis, Parra *et al*. (2010) obtained different patterns of community structure for hummingbirds (Trochilidae) and their separate subclades (emeralds, mangoes, brilliants) (Parra *et al*., 2010). Chalmandrier *et al*. (2013) manipulated phylogenetic extent through randomization (within-clades and between-clades) to uncover the effects of biotic interactions, which were masked by environmental filtering at large phylogenetic extents.

There are several promising avenues for further integration of phylogenetic scale into community ecology. First, the use of multiple metrics covering a range of phylogenetic scales should become the standard practice to provide more complete information about community structure. Even though cross-grain and cross-extent approaches can be informative, as illustrated by the case studies above (Swenson, 2009; Parra *et al*., 2010; Chalmandrier *et al*., 2013; Parmentier *et al*., 2014; Mazel *et al*., 2016), phylogenetic grain and extent might prove hard to manipulate separately, as changes in one often produce changes in the other (e.g. an increase in the phylogenetic extent increases also the grain captured by NRI and NTI) (Webb *et al*., 2002; Swenson, 2009). Second, experiments can be designed to target specific phylogenetic scales, where the processes of competition and environmental filtering have been inferred to operate (Godoy *et al*., 2014). Third, the grain of the analysis might be extended to include within-species processes, relevant to community structure (e.g. trait variation, demographic structure), as advocated by the field of community genetics (Hersch-Green *et al*., 2011).

Phylogenetic scale might produce further insights for invasion ecology and conservation. Darwin’s naturalization hypothesis postulates that communities are more open to invasion by unrelated species, while close relatives of the resident species are less likely to invade (reviewed by Thuiller *et al*., 2010), which makes the susceptibility of a community toward invasion phylogenetic scale-dependent (Thuiller *et al*., 2010; Godoy *et al*., 2014). Moreover, phylogenetic metrics that target different phylogenetic depths might help to guide conservation priorities (Redding *et al*., 2014). Some communities consist of closely related species that show a high degree of phylogenetic and functional redundancy (e.g. mammals of South America), while others include a variety of species with unique evolutionary histories (e.g. mammals of Africa) (Oliveira *et al*., 2016), which might qualify these communities to receive heightened attention from conservation biologists (Redding *et al*., 2014).

### Biogeography and niche conservatism

Biogeographic patterns, such as species distributions and diversity gradients, are largely shaped by the conservatism of ecological niches (Wiens & Graham, 2005), and much literature has been dedicated to the question whether or not niches are conserved (Wiens & Graham, 2005; Pyron *et al*., 2015). Yet, it might be more fruitful to reframe this question in terms of phylogenetic scale. In particular, niches might be conserved over some phylogenetic scales, but not others, such that we can investigate how the scale-dependence of niche conservatism contributes to various biogeographic patterns.

Taxa often show different patterns of regional richness, which might partly result from the fact that their climatic niches are conserved over different phylogenetic scales (Wiens & Graham, 2005; Donoghue, 2008; Buckley *et al*., 2010). For example, many clades of mammals have generated high richness in the tropics but never invaded the temperate, presumably because their climatic niches have been highly conserved over large phylogenetic scales. Relaxed conservatism in the clade of rabbits and hares, in contrast, might have allowed the group to colonize and diversify in temperate climates (North America, Eurasia) (Buckley *et al*., 2010). Shifts in the climatic niche, which typically span only several nodes in the phylogeny and a short period in the history of a clade, might lead to diversification episodes that enrich the diversity of regional biotas (Wiens & Graham, 2005; Donoghue, 2008; Buckley *et al*., 2010; Glor, 2010). Consequently, the evaluation of niche conservatism across phylogenetic scales might inform us about the formation of diversity gradients.

Diversity patterns may be further influenced by the effects of niche conservatism on regional extinctions (Jackson & Weng, 1999; Cahill *et al*., 2012). For example, many genera of trees were wiped out by climatic changes during Pleistocene (Jackson & Weng, 1999). Yet, the same changes exterminated only a few tree families, presumably because the climatic niches of trees are more conserved at the genus-level than at the family-level (Donoghue, 2008; Cahill *et al*., 2012). The extinction footprint of climate change might therefore depend on the phylogenetic scale of niche conservatism. Evaluating scale-dependent vulnerability to extinction seems particularly relevant in the face of the on-going climatic and land-use changes, and the results of such an evaluation might inform us about the patterns of loss of phylogenetic diversity (Purvis *et al*., 2000). Taken together, even though it has long been recognized that niches are conserved to varying degrees (Blomberg *et al*., 2003; Wiens & Graham, 2005; Donoghue, 2008; Buckley *et al*., 2010; Price *et al*., 2014; Pyron *et al*., 2015), few studies have systematically investigated this variation across various phylogenic scales despite the promise that such an investigation might enhance our understanding of diversity gradients.

### Macroecology

Macroecologists are concerned with statistical patterns observed across large spatial and temporal scales, such as the latitudinal diversity gradient, body size distributions, species-area relationships or species-abundance distributions (Brown, 1995). Phylogenetic scale is rarely explicitly considered in macroecological research. Yet, by exploring the patterns across different phylogenetic scales, it is possible to identify which patterns are truly universal and which are not because they disintegrate over certain phylogenetic scales (Marquet *et al*., 2004; Storch & Šizling, 2008). This exploration might reveal the mechanisms that produced these patterns (Marquet *et al*., 2004; Storch & Šizling, 2008).

Many of the well-known macroecological patterns emerge only across a narrow range of phylogenetic scales. For example, the decrease in regional richness with latitude (latitudinal diversity gradient) holds for higher taxa (mammals, birds, or plants) but often breaks down across small phylogenetic scales (e.g. within penguins, hares, aphids, ichneumonids, Proteacea) (e.g. Kindlman *et al*., 2007; Buckley *et al*., 2010) (Fig. 1a). Likewise, species abundance and body mass are negatively correlated across birds and mammals (Damuth, 1981; Isaac *et al*., 2011) (Fig. 1b), but the correlation sometimes disappears across narrowly defined taxa, and may even become positive in select tribes of birds (Cotgreave, 1994). Small-bodied species can be highly abundant, presumably because their metabolic requirements are low and that raises the carrying capacities of their populations. However, over small phylogenetic scales, which comprise ecologically similar and potentially competing species, large-bodied species are competitively superior and consequently more abundant than small-bodied species, which reverses the directionality of the correlation between body size and population abundance (as illustrated in Fig. 1d) (Cotgreave, 1994).

Moreover, the exploration of macroecological patterns across phylogenetic scales may inform us about their presumed universality. Specifically, the species-area relationship (SAR) and species-abundance distribution (SAD) were believed to universally conform to particular mathematical forms (the power-law function and the lognormal distribution, respectively) (Preston, 1948; Rosenzweig, 1995). However, if two sister taxa follow power-law SARs and lognormal SADs that differ in their parameters, it can be demonstrated mathematically that the clade containing both sister taxa cannot follow either the power-law SAR or the lognormal SAD (Storch & Šizling, 2008; Šizling *et al*., 2011). Therefore, even though some macroecological patterns have been considered classic examples of ecological laws, cross-scale analyses often reveal that these patterns cannot be truly universal. Moreover, the fact that some patterns disintegrate across phylogenetic scales implies either that the theories to explain these patterns are fundamentally ill-founded, or that the patterns pertain to certain phylogenetic scales only (Storch & Šizling, 2008). The latter possibility suggests that phylogenetic scales form phylogenetic domains within which our theories apply; an explicit delimitation of these domains might then further inform the theory.

### Phylogenetic scale in practice

The above overview demonstrates that some fields have considered the concept of phylogenetic scale more extensively and systematically than others and that the approach used by different fields often varies in terms of vocabulary (e.g. tree depth, taxonomic rank, clade age) as well as methodology (e.g. varying phylogenetic extent, grain, tree-depth). Unifying some of these efforts under the concept of phylogenetic scale might provide the common ground for cross-field discussion. Previous research often investigated phylogenetic scale within an exploratory framework, whereby patterns were identified across a range of scales and then explained in the light of specific events or mechanisms. Another possible approach relies on testing a priori hypotheses, which are based on mechanisms assumed to operate over certain phylogenetic scales. Both of these approaches (exploratory and hypothesis-testing) have their strengths, and either may be appropriate, depending on the objective of a given study.

In this section, we consider the conceptual and methodological approaches to investigate patterns across phylogenetic scales. Above, we illustrated that many attributes of interest vary across scales (e.g. diversification rate, niche conservatism, geographic distribution, statistical relationships). These attributes may vary randomly or systematically across the phylogeny, be more prevalent at particular scales, or stay unchanged across a discrete set of mutually nested clades (Box 2). We refer to the latter as a domain of phylogenetic scale which, in analogy to spatial domains (Wiens, 1989), corresponds to a segment of the phylogeny that reveals homogeneity in the attribute of interest. Below, we elaborate on each of these concepts together with the methods for their empirical evaluation.

### Choice of phylogenetic scale

While most researchers are aware that the choice of scale can influence inferences about patterns or processes, all too often the choice of scale, be it spatial, temporal or phylogenetic, is influenced by data availability or other logistical concerns. However, this might lead to a mismatch between the phylogenetic scale of analysis and the biological question addressed. For example, to test the hypothesis that competition limits species coexistence locally, small phylogenetic extents (e.g. genera, or clades where species can be reasonably assumed to compete with each other) should be preferred to large extents where most species are highly unlikely to compete (e.g. the entire classes, such as birds and mammals). However, even with a specific question at hand, it can be hard to choose a single most suitable phylogenetic scale, and working across multiple scales might be more appropriate.

#### BOX 2: Research across phylogenetic scales

Many attributes, such as diversification rate, niche conservatism, or community structure, vary across phylogenetic scales (Table 1). They can vary in three different ways:

**(a) Scale-dependence** refers to the situation when the attribute of interest changes across phylogenetic scales without any clear trend. In this case, the results from one scale might be difficult to extrapolate to another scale.

**(b) Scaling** occurs when the attribute of interest varies systematically along the scale axis. The interpretation of scaling is at least threefold, depending on the underlying mechanism (note that only one of the interpretations is biological):

1. **Statistical scaling** is a sample-size effect whereby the statistical power of the analysis increases with grain. Consequently, the attribute under study appears to change systematically from small to large scales (Machac *et al*., 2013). While the inferred values of the attribute itself may be technically correct, their systematic variation across scales is biologically irrelevant.
2. **Methodological artifacts** result when a statistical analysis becomes increasingly misleading toward the deep nodes of the phylogeny (i.e. large scales), resulting in incorrect and potentially biased estimates for the attribute of interest (e.g. ancestral reconstructions under dispersal-vicariance models tend to suggest that the ancestor occupied all of the regions examined) (Ronquist, 1997). Methodological artifacts can be mitigated using various statistical corrections or when the results are validated using supplementary data, such as fossils.
3. **Phylogenetic scaling in the strict sense** occurs when the studied attribute changes across scales because the underlying biological process changes. True scaling can therefore inform us about the processes which generate the patterns observed across scales. If the scaling can be described mathematically, it may allow extrapolation across scales, even those not included in the original study, i.e. *downscale* or *upscale* the patterns under study.

**(c) Domains of scale** refer to the segments of the phylogeny (e.g. clades or tree depth) within which the attribute of interest appears relatively unchanged. The attribute might change abruptly between domains, indicating changes in the underlying biological processes. Therefore, it is possible to extrapolate across phylogenetic scales within, but not between, domains.

**Figure:** Numerous variables can be studied across phylogenetic scales. These may include 436 diversification rates, statistical relationships between ecological variables, parameters of statistical 437 distributions, community metrics, or niche conservatism (see Table 1). Phylogenetic scale may be 438 defined in terms of clade age, clade size, tree depth, degree of phenotypic divergence, etc. (Box 1).

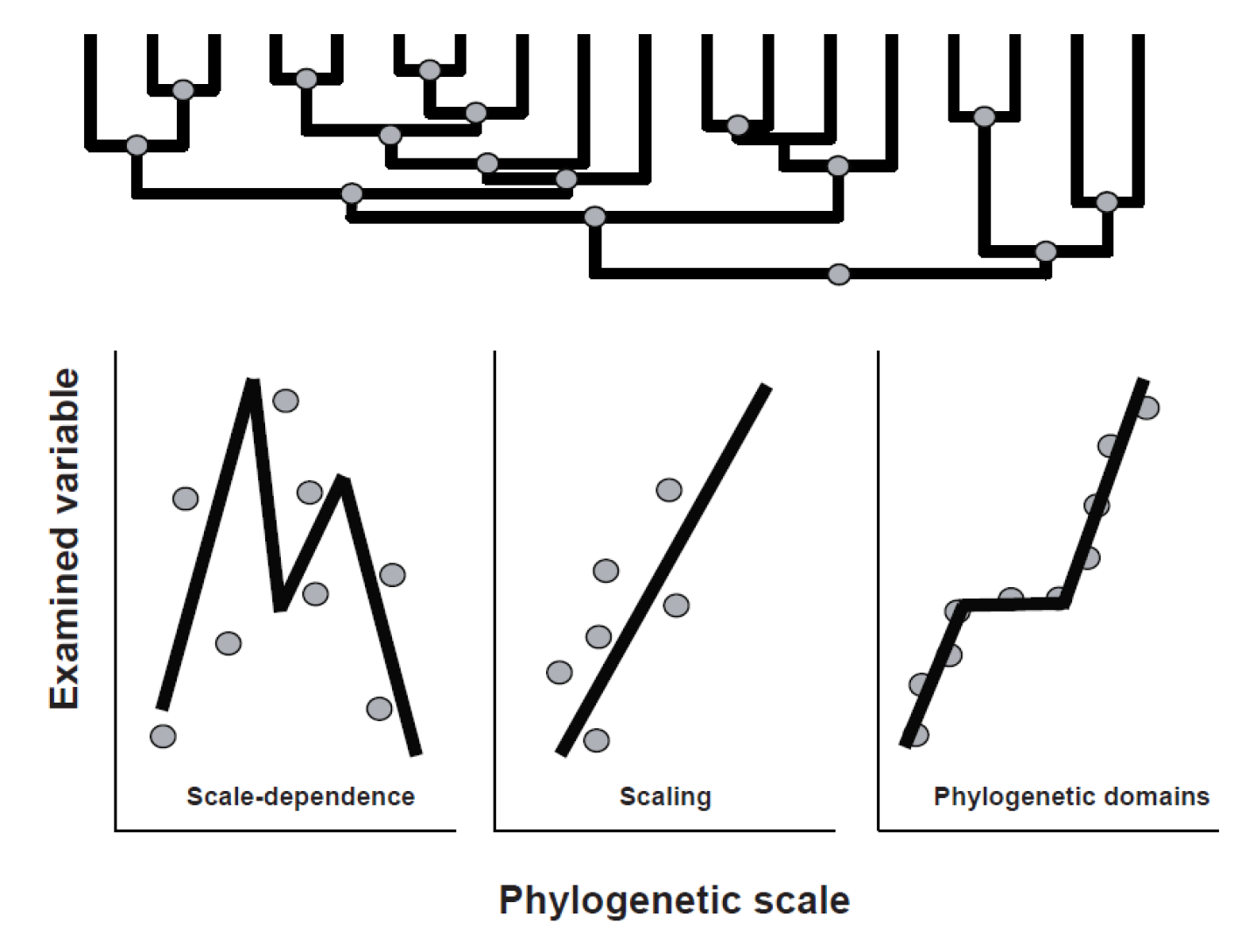

**Table 1.** Ecological and evolutionary attributes that can vary across phylogenetic scales. Each attribute is listed together with examples of methods for its evaluation.

| Field | Examined attribute | Methods for the evaluation of the attribute | Implementation in R |
| --- | --- | --- | --- |
| **Evolution and diversification** | diversification mode | coalescent inference to distinguish between accelerations, slowdowns, and saturation (Morlon *et al.*, 2010) | RPANDA (Morlon 2016) |
|  | diversification rate | product-moment estimators (Magallon & Sanderson, 2001), equal-splits measures (Jetz *et al.*, 2012) | ape (Paradis *et al.*, 2004), geiger (Harmon *et al.*, 2008) |
|  | slowdown strength | gamma statistic (Pybus & Harvey, 2000) | laser (Rabosky and Schliep 2013) |
| **Community ecology** | community structure and phylogenetic diversity | phylometrics (NRI, NTI, MNND, MPD, PD) (Faith, 1992; Webb *et al.*, 2002; Swenson, 2009) | picante (Kembel *et al.*, 2010), PhyloMeasures (Tsirogiannis and Sandel 2017) |
| **Niche conservatism and trait evolution** | phylogenetic signal | Pagel's lambda (Freckleton *et al.*, 2002), Blomberg's K (Blomberg *et al.*, 2003) (but see Revell *et al.*, 2008) | geiger (Harmon *et al.*, 2008), picante (Kembel *et al.*, 2010) |
|  | evolutionary rates | Brownian motion model (Edwards & Cavalli-Sforza, 1964; Felsenstein, 1985), Ornstein-Uhlebeck model (Hansen, 1997), ACDC model (Blomberg *et al.*, 2003) | ape (Paradis *et al.*, 2004), geiger (Harmon *et al.*, 2008) |
| **Biogeography and macroecology** | statistical relationship | linear, polynomial, exponential, lognormal functions, etc. | base (R Core Team 2017), nlme (Pinheiro *et al.* 2013) |
|  | relationship strength | Pearson's correlation, Spearman's correlation | base (R Core Team 2017) |

**Table 2.** Methods that work across phylogenetic scales. These methods return comprehensive results for the entire phylogeny, which can be used to investigate scale-dependence, phylogenetic scaling, and the domain of scale (Box 2). The results of each method are briefly explained, together with further references and the software for its empirical implementation.

| Studied pattern/process | Method | Results | Software and references |
| --- | --- | --- | --- |
| **Diversification** | BAMM, MEDUSA, REVBayes | shifts in diversification rates and regimes (constant diversification, accelerations, slowdowns) across the entire phylogeny | BAMMtools (Rabosky, 2014), geiger (Harmon *et al.*, 2008; Alfaro *et al.*, 2009), REVBayes (Hohna *et al.*, 2016) |
| **Trait evolution** | BAMM, SURFACE, NODIV, MOTMOT, PIC, OU, ACDC, BM | changes in the values and rates of traits (morphological, behavioral, physiological, molecular) across the phylogeny | BAMMtools (Rabosky, 2014), surface (Ingram & Mahler, 2013), nodiv (Borregaard *et al.*, 2014), MOTMOT (Thomas & Freckleton, 2012), ape and geiger (Edwards & Cavalli-Sforza, 1964; Felsenstein, 1985; Hansen, 1997; Blomberg *et al.*, 2003; Butler & King, 2004; Paradis *et al.*, 2004; Harmon *et al.*, 2008) |
| **Geographic distributions** | BIOGEOBEARS, LAGRANGE, NODIV, DIVA | dispersal and colonization events, shifts in the geographic distributions | BioGeoBEARS (Matzke, 2014), LAGRANGE (Ree *et al.*, 2008), nodiv (Borregaard *et al.*, 2014), DIVA (Ronquist, 1997; Ronquist & Sanmartin, 2011) |

### Multiple phylogenetic scales

In some cases, it might be illuminating to study how an attribute of interest behaves at multiple phylogenetic scales. This strategy seems particularly relevant in large phylogenies whose numerous clades may show very different patterns in the attribute of interest (e.g. diversification rate, the strength of niche conservatism, patterns of community phylogenetic structure) (Fig. 1). For example, Cetacean systematists had long been perplexed as to why there is little correspondence between diversification dynamics estimated from the fossil record and phylogenetic trees (Morlon *et al*., 2011). The correspondence between the two datasets emerged only when considering diversification heterogeneity across clades. The results suggested that individual clades, which represented small phylogenetic extents (i.e. ronquals, ocean dolphins, porpoises and beaked whales), had their own dynamics that were obscured at the phylogenetic extent of Cetaceans as a whole (Morlon *et al*., 2011).

Several methods have been developed to investigate patterns across phylogenetic scales (Table 1, Table 2). The method introduced by Borregaard et al. (2014), for example, proceeds across the phylogeny and identifies the nodes whose descendant clades underwent conspicuous geographic, phenotypic, or ecological shifts (Borregaard *et al*., 2014). Similar methods have also been developed to investigate community structure across various phylogenetic grains (Parmentier *et al*., 2014) and phylogenetic extents (Chalmandrier *et al*., 2013). In macroevolution, statistical algorithms that proceed across the entire phylogeny are not uncommon and have been used to identify shifts in diversification rates (e.g. BAMM, MEDUSA, REVBAYES) (Alfaro *et al*., 2009; Rabosky, 2014; Hohna *et al*., 2016). These shifts then delimit those segments of the phylogeny whose diversification is relatively homogeneous, such that the segments might be perceived as phylogenetic grains for further analysis (= elementary and homogeneous unit of analysis) or as phylogenetic domains (see Box 2). Phylogenetically independent contrasts (Felsenstein, 1985) are also calculated for the entire phylogeny and thus capture trends across an inclusive range of phylogenetic scales. Yet, they are rarely explored with respect to the phylogenetic scale itself (e.g. contrasts might decline from the root toward the tips, indicating progressively decreasing evolvability in the trait of interest). Instead, most studies use contrasts to confirm an overall correlation between two variables (Felsenstein, 1985). These examples together illustrate the range of tools that can be readily used for cross-scale analyses. Nonetheless, many studies work with select clades only, despite the commonly cited concern that clade selection is typically non-random and might bias the results (Phillimore & Price, 2008), while cross-scale analyses remain relatively underutilized.

Two potential issues, associated with the evaluation of all nodes within a phylogeny, are data non-independence and nestedness. Non-independence can be readily accommodated by widely used comparative methods (e.g. PIC, PGLS) (Felsenstein, 1985; Freckleton *et al*., 2002). These methods typically estimate the same parameters as their conventional counterparts (e.g. intercepts, regression slopes, group means) but adjust the confidence intervals of these parameters based on the inferred degree of phylogenetic correlation in the data (Freckleton *et al*., 2002). The nestedness of the data is more difficult to accommodate. For example, the diversification rate of a clade is inherently determined by the rate values across its constituent subclades. Nestedness therefore extends beyond the phylogenetic correlation of rate values and reflects how the value for a clade is produced by the subclade values. This information cannot be readily accommodated under the currently available comparative methods, such that phylogenetic corrections do not guarantee proper estimates of statistical significance across nested data. For these reasons, we argue that parameter estimates can be extracted, compared, and analyzed across nested clades, but their significance needs to be interpreted cautiously. New theory that would illuminate how different attributes of interest (e.g. diversification rates, regression slopes, phylogenetic signal) combine and compound across nested hierarchies, as well as the methods to capture these correlations, might be needed.

### Phylogenetic scaling

Scaling is a trend in the attribute of interest across the phylogenetic hierarchy (e.g. across clades ordered with respect to clade age, taxonomic rank, phylogenetic depth, or some other suitable measure of phylogenetic scale) (Box 1). For example, if scale is defined in terms of clade age, then scaling in niche conservatism is a systematic trend in conservatism with clade age.

Phylogenetic scaling should be most prevalent across mutually nested clades because the patterns associated with larger clades are determined by the patterns of clades nested within them (or vice versa). For example, diversification rate of a clade is determined by the rate values of its subclades, similarly as species richness of a spatial plot is determined by the richness of its subplots. Consequently, it should be possible to predict the value of an attribute (e.g. diversification rate, regression slopes, phylogenetic signal) at a particular phylogenetic scale from the knowledge of those values across other scales, much like it is possible to estimate species richness within large geographic areas, based on the knowledge of richness within small areas (Storch, 2016). When characterized mathematically, phylogenetic scaling should allow for predictions across phylogenetic scales not covered by the phylogeny under consideration (i.e. upscaling or downscaling).

Statistical methods of cross-scale analysis (see above) are particularly suitable to investigate phylogenetic scaling (e.g. BAMM, MEDUSA, PIC) (Felsenstein, 1985; Alfaro *et al*., 2009; Pavoine *et al*., 2010; Chalmandrier *et al*., 2013; Borregaard *et al*., 2014; Parmentier *et al*., 2014; Rabosky, 2014) (Table 1, Table 2). These methods explore macroevolutionary patterns across the entire phylogeny (e.g. changes in the rates of speciation and extinction) (Alfaro *et al*., 2009; Ingram & Mahler, 2013; Rabosky, 2014), analyze multiple grains within the same extent (Pavoine *et al*., 2010; Borregaard *et al*., 2014; Parmentier *et al*., 2014) or change the extent while holding the grain unchanged (Parra *et al*., 2010; Chalmandrier *et al*., 2013), thus producing a range of results for different phylogenetic scales. These results can then be further investigated specifically with respect to the scale axis (e.g. diversification rates decline over time, competition increases with species relatedness toward the shallow nodes of the phylogeny, etc.), as detailed in Box 2.

### Domains of phylogenetic scale

When moving along the scale axis, the values of an attribute might sometimes change abruptly. Such discontinuities provide the opportunity to delimit domains of phylogenetic scale (Box 2). Domains are discrete segments of a phylogeny, such as monophyletic clades, taxonomic ranks, sets of nodes, or tree depth, which show homogeneity in the attribute of interest (i.e. diversification rate, statistical correlation, or phylogenetic signal). By definition, the attribute does not vary substantially within a domain, but changes between domains. Phylogenetic domains may therefore provide insights into the processes which operate over different segments of a phylogenetic tree.

Traditionally, phylogenetic domains were delimited by taxonomists whose objective was to organize species into biologically meaningful units, such as families, orders, or classes. These units are based mostly on morphological and ecological attributes. Phylogenetic domains, however, can also consist of clades that show diversification homogeneity, comparable rates of morphological evolution, or similar life-history trade-offs. Therefore, the delimitation of the domains might rely on the natural history of the group (key innovations, episodes of historical dispersal, extinction events, etc.) but also on statistical methods that do not require any such prior knowledge (Table 2). The statistical delimitation might be more transparent and reproducible, but the resulting domains may be harder to interpret biologically. Still, domains delimited statistically may reveal otherwise unnoticed evolutionary events and relevant shifts in a clade’s history, which may have contributed to its extant diversity.

Phylogenetic domains may also facilitate statistical inference, given that most comparative methods assume that the attributes analyzed are homogeneous (e.g. regression slopes do not vary across genera within the analyzed family, diversification is homogenous across the analyzed lineages) and return spurious results when applied to clades that show a mixture of patterns and processes (Morlon *et al*., 2011; O’Meara, 2012) (Fig. 1d). Phylogenetic domains may therefore help to identify when comparative methods report reasonably reliable results and when their conclusions must be interpreted with caution because the results span different domains and the underlying assumptions have been violated.

## Conclusion

It is well established that different processes dominate over different spatial and temporal scales. Phylogenetic scale, however, has received limited attention although much research in ecology and evolution today relies on molecular phylogenies (Table 1, Table 2). Explicit consideration of different aspects of phylogenetic scale, including grain, extent, scale dependence, phylogenetic scaling, and the domains of phylogenetic scale can therefore inform multiple fields (e.g. macroevolution, community ecology, biogeography and macroecology).

We discussed phylogenetic scale largely in isolation from spatial and temporal scales, but these types of scale will often be related. For instance, competitive exclusion may be prominent among closely related species within local communities over short time periods (Cavender-Bares *et al*., 2009). In contrast, plate tectonics might influence deeper nodes in a phylogeny and operate over broad geographic extents (Willis & Whittaker, 2002). In some notable cases, however, the spatial and phylogenetic scales may not be related. Diversity anomalies, such as New Caledonia or Madagascar, represent examples of decoupling where rich biotas that encompass extensive phylogenetic scales diversified in a relatively small region (Espeland & Murienne, 2011). In contrast, recent radiations within grasses and rodents have had a large geographic footprint but encompass only a few relatively young clades (Edwards *et al*., 2010). Evaluating when different types of scale are coupled (or decoupled) may yield new insights into the evolutionary history of different clades and regions (Wiens, 1989; Rosenzweig, 1995; Willis & Whittaker, 2002).

We hope that the perspective presented here will spur further theoretical, empirical, and methodological research. Explicit consideration of phylogenetic scale may turn our focus away from particular mechanisms toward the appreciation of the interplay of multiple processes that together, but over different phylogenetic scales, shape the diversity of life.

## Acknowledgements

Funding was provided by the NSF program Dimensions of Biodiversity (DEB-1136586) and by the Grant Agency of the Czech Republic (14-36098G).

